# Lytic bacteriophage have diverse indirect effects in a synthetic cross-feeding community

**DOI:** 10.1101/560037

**Authors:** Lisa Fazzino, Jeremy Anisman, Jeremy M. Chacón, Richard H. Heineman, William R. Harcombe

## Abstract

Bacteriophage shape the composition and function of microbial communities. Yet, it remains difficult to predict the effect of phage on microbial interactions. Specifically, little is known about how phage influence mutualisms in networks of cross-feeding bacteria. We modeled the impacts of phage in a synthetic microbial community in which *Escherichia coli* and *Salmonella enterica* exchange essential metabolites. In this model, phage attack of either species was sufficient to inhibit both members of the mutualism; however, the evolution of phage resistance ultimately allowed both species to attain yields similar to those observed in the absence of phage. In laboratory experiments, attack of *S. enterica* with P22*vir* phage followed these modeling expectations of delayed community growth with little change in the final yield of bacteria. In contrast, when *E. coli* was attacked with T7 phage, *S. enterica*, the non-host species, reached higher yields compared to no-phage controls. T7 increased non-host yield by releasing consumable cell debris and by driving evolution of phage resistant *E. coli* that secreted more carbon. Additionally, *E. coli* evolved only partial resistance, increasing the total amount of lysed cells available for *S. enterica* to consume. Our results demonstrate that phage can have extensive indirect effects in microbial communities, and that the nature of these indirect effects depends on metabolic and evolutionary mechanisms.

## Introduction

Bacteriophage significantly influence microbial community structure and function [1]. Phage limit the size of bacterial populations, which can change microbial community composition. For example, phage kill >20% of marine bacteria every day [2]. Viral infection of bacterial populations not only impacts the composition of bacterial communities, but also influences emergent community functions such as the rate at which nutrients are converted into biomass [3]. As a result, phage critically influence biogeochemical cycling [4], biotechnology [5], the food industry [6], and human health [7,8]. Despite the importance of phage in microbial communities, we cannot reliably predict the impact of phage on the composition and function of communities. As we strive to manage microbial communities, we must improve our understanding of phage effects in multi-species systems.

Phage alter bacterial communities by changing the abundance of competing species. When phage kill dominant competitors, weaker competitors that are not susceptible to the phage can flourish [9]. These opposing changes in species abundance alter species ratios [10–12]. Sometimes, species ratios rapidly revert to pre-phage frequencies once a host evolves phage resistance, but costs of resistance can generate persistent changes in community composition following phage addition [10]. In contrast to the volatility of species ratios, phage attack of one species often has little impact on total community biomass. In communities of competitors, the reduction in host biomass is compensated for by the growth of non-host competitors [13]. Taken together, in competitive systems, phage alter species ratios, but have little impact on total microbial biomass.

Much less is understood about how phage influence cooperative networks in microbial communities. Microbial communities are often organized into cross-feeding webs in which each species relies on metabolites excreted by others [14,15]. Networks of metabolic dependencies have been described in marine, terrestrial, and human-associated microbial communities [14]. While phage are likely present in all of these systems, the impact of phage on the composition and function of cross-feeding microbial communities remains under-studied. However, responses of cross-feeding communities to abiotic disturbances can inform the impact that phage may have. Obligate mutualism tends to constrain species ratios such that communities will converge to an equilibrium frequency from any initial composition [16,17]. This constraint on species ratios means that limiting one species should indirectly inhibit cross-feeding partners, thereby decreasing total community biomass [18]. For example, antibiotic treatment that inhibited the most sensitive member of an obligate cross-feeding community was sufficient to inhibit growth of the entire community [19]. We therefore predict that phage attack on one member of a cooperative network is likely to limit growth of the entire microbial community by depriving obligate cross-feeding species of nutrients but will not affect species ratios.

However, the hypothesis that phage will alter community biomass but not composition has several underlying assumptions that may not hold. First, it assumes that bacteria only obtain nutrients from the excretions of bacterial partners. Yet, there is a rich body of literature suggesting that phage-mediated cell lysis releases nutrients into the environment. Indeed, this ‘viral shunt’ is thought to play a major role in global nutrient cycling [20–22]. Furthermore, release of additional nutrients may eliminate the obligate nature between bacterial species and reduce constraints on species ratios [23,24]. Second, the hypothesis overlooks possible ecological consequences of the evolution of phage resistance. For example, it assumes that phage resistance does not alter the exchange of cross-fed nutrients. If phage resistance causes changes in either nutrient secretion or uptake, it could radically alter species ratios. Phage are likely to dramatically impact cross-feeding networks; however, it remains difficult to predict the impacts.

In this study, we sought to determine the effects of phage attack on cooperative communities by combining resource-explicit mathematical modeling and wet-lab experiments of a synthetic cross-feeding co-culture composed of *Escherichia coli* and *Salmonella enterica* [17,25]. An *E. coli* strain auxotrophic for methionine was paired with an *S. enterica* strain that was evolved to secrete methionine [25]. The pair forms an obligate mutualism in lactose minimal medium as *S. enterica* cannot consume lactose and instead relies on acetate excreted by *E. coli*. To this community, we added either an *E. coli*-specific (T7) or *S. enterica-specific* (P22*vir*) lytic phage and tracked community responses (Fig 1a). As a null hypothesis, we predicted that cross-feeding would constrain species ratios and therefore phage attack of one species would inhibit the entire community. However, we anticipated that phage resistance would evolve, making biomass reduction temporary. We found that both phage delayed community growth, but T7 infection of *E. coli* led to surprising changes in species ratios.

**Figure 1.**
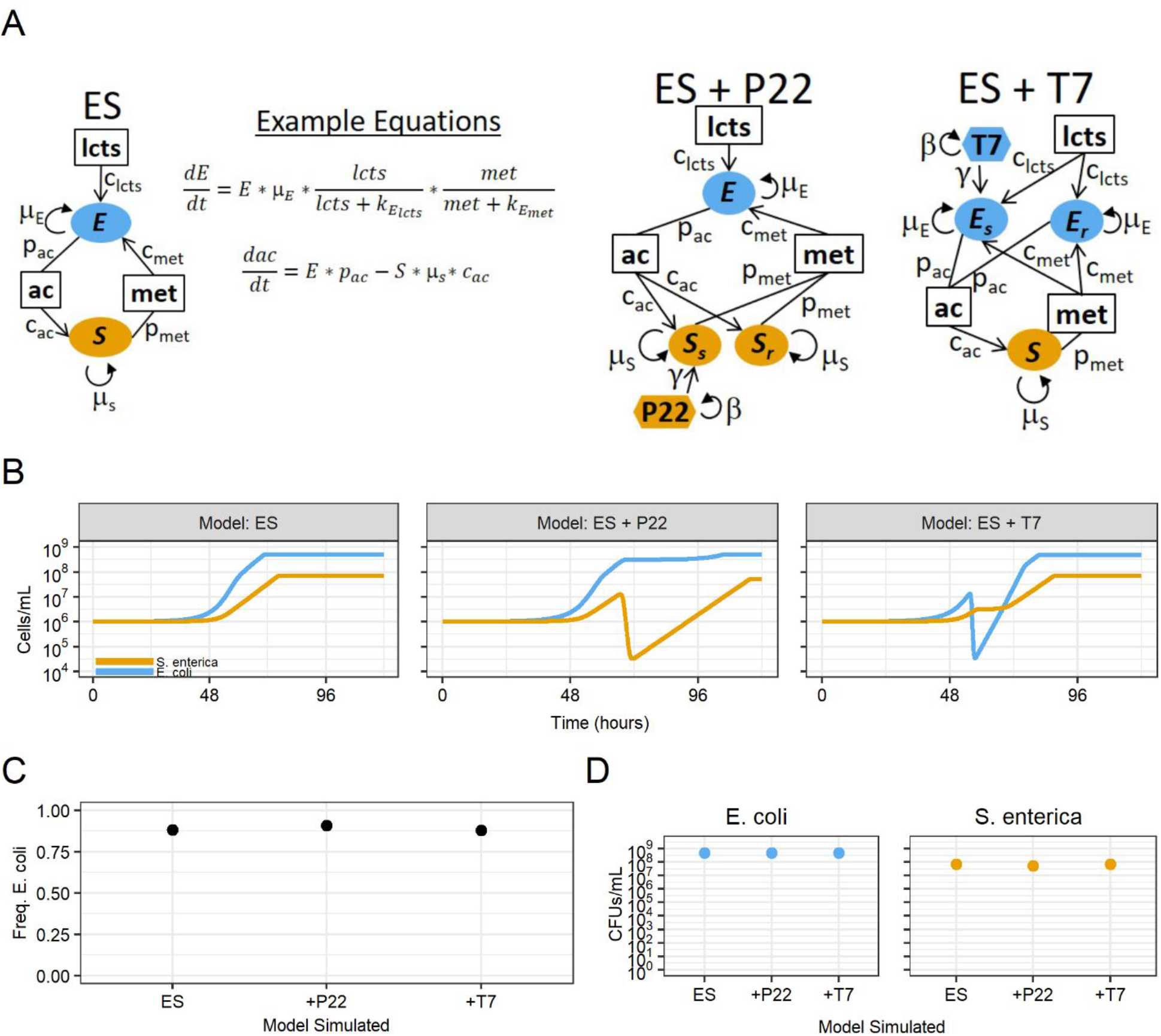
Mathematical model predicting consequences of phage infection of cooperative co-cultures. **A)** Schematic of models for systems with *E. coli* (E), *S. enterica* (S) and phage (T7 or P22). Bacteria are represented by ovals. Phage sensitivity (s) or resistance (r) is indicated by subscripts. Phage are indicated by hexagons and are colored to match their bacterial host. Boxes indicate metabolites (lcts = lactose, ac = acetate, met = methionine). Parameters are next to interaction arrows. c_x_ = consumption rate of subscript nutrient, p_x_ = production rate of subscript nutrient, μ = growth rate (h^−1^), β = burst size, γ = adsorption constant. **B)** Simulated growth curves with and without phage treatment. Yellow = *S. enterica*, blue = *E. coli*. **C)** Species ratios represented with frequency of *E. coli* at time = 125 hours. **D)** Final densities of bacteria (cells/mL) in mathematical models at time = 125 hours.

## Results

### Resource-explicit model suggests phage have little impact on final community composition

We used a resource-explicit model to predict how a cooperatively growing bipartite bacterial community would respond to lytic phage attack under batch growth. We modeled densities of *E. coli* (E), *S. enterica* (S), and phage (T7 or P22*vir*), as well as the concentrations of lactose, methionine, and acetate (Fig 1a, Supplemental Methods). Growth of each bacterial species was a function of maximum growth rate (μ_max_) and Michaelis-Menten saturation parameters (k_m_) for essential metabolites with multiplicative growth limitation when species are limited by multiple metabolites. The model was scaled to individual bacterial cells. Bacterial death due to phage infection was modeled as a linear interaction between phage and host, and modified by an adsorption (i.e. predation) constant (γ). Phage attack generated new phage particles at a rate set by the burst size (β). Phage-resistant hosts (E_R_ or S_R_) were initiated at a frequency of 0.1% in each bacterial population to create variation allowing for the spread of resistance alleles following phage introduction. The growth parameters (μ_max_ and k_m_) were equal for sensitive and resistant bacteria, resulting in no cost of resistance. Acetate and methionine were produced (p) and consumed (c) at a rate proportional to cell growth. Parameters were informed by literature and adjusted to match experimental observations in the absence of phage (Supp Table 1).

In the absence of phage, the community exhibited expected dynamics. *E. coli* and *S. enterica* cross-fed and grew until all of the lactose was consumed. *E. coli* reached a carrying capacity of 5.1 × 10^8^ cells/mL, and *S. enterica* reached a carrying capacity of 6.8 × 10^7^ cells/mL if both species were present. In absence of either partner, no growth was observed. The community converged to 88% *E. coli* regardless of the ratio at which bacteria were started, consistent with previous observations (Fig 1c, Supp Fig 1)[17]. In the absence of phage, resistant bacterial genotypes did not increase in percentage of the host population, but were not eliminated (Supp Fig 2).

The model predicted that the presence of phage increases the time required for the community to reach carrying capacity, but has little impact on the final species yields. Both T7 and P22*vir* rapidly killed all sensitive hosts (Fig 1b, Supp Fig 2). The reduction in the host population reduced the amount of cross-feeding, thereby temporarily stalling community growth. However, phage-resistant hosts rapidly increased in abundance, allowing the community to reach carrying capacity. Host species reached slightly lower final densities as a result of phage attack (Fig 1d). This reduction is because sensitive host cells consume resources before they are killed by phage, and fewer resources are therefore available for growth of the resistant host. However, the final yield of the host is only reduced ~1.03 – 1.32 fold (by 4-25%) because sensitive hosts are killed before they are able to consume many resources. No change was observed in the final abundance of the non-host bacteria.

### P22*vir* followed expectations but T7 altered bacterial yields in wet-lab experiments

Using our experimental cross-feeding system in lactose minimal media, we tested how the bacterial community responded to *S. enterica*-specific P22*vir* or *E. coli*-specific T7 lytic phage attack. Five replicates and five controls were grown for each phage treatment. Communities were started with a multiplicity of infection (MOI) of 0.01, and bacterial populations of 5×10^5^ cells/mL per species (Supp Table 2). Resistant host cells arose via mutation during cooperative community growth; they were not seeded into the host population. Growth of each bacterial species was tracked with a unique fluorescent marker which could be converted to a species-specific OD (Fig 2a, see methods). After growth, co-cultures were plated for *E. coli* and *S. enterica* colony forming units (CFUs), and T7 or P22*vir* plaque-forming units (PFUs).

**Figure 2.**
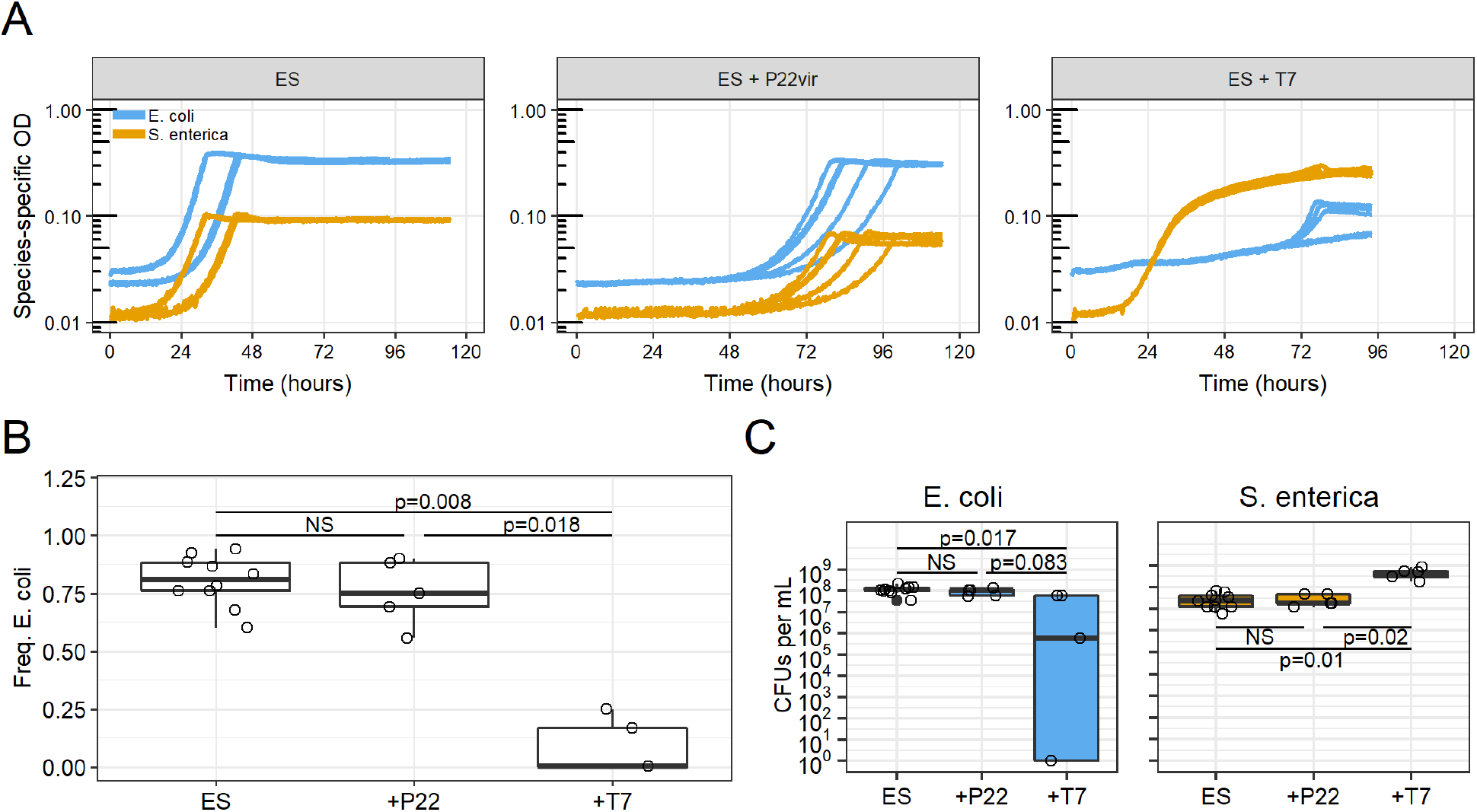
Experimental data of cooperative co-culture growth and the effect of adding phage. **A)** Species-specific growth curves calculated by transforming fluorescence into species-specific OD for three treatments: no phage (ES), *S. enterica*-specific P22*vir* phage (ES + P22), or *E. coli*-specific T7 phage (ES + T7). *E. coli* (CFP) = blue, *S. enterica* (YFP) = yellow. **B)** Species ratios represented with frequency of *E. coli* at the end of growth. Ratios were calculated from plated CFUs. **C)** Boxplot depicting measured CFUs of *E. coli* or *S. enterica* at the end of growth. Statistical significance determined with Mann-Whitney-U test with FDR multiple hypothesis correction.

Attack of *S. enterica* with P22*vir* delayed community growth as compared to the no-phage control (p = 0.0037, Supp Fig 3) but had little impact on final community composition, as predicted by our model. The final frequency of *E. coli* was not different from communities with no phage (p = 0.513, Fig 2b). Furthermore, neither *S. enterica* (p=0.95) nor *E. coli* (p = 0.35) final densities were significantly changed by the addition of P22*vir* to the cross-feeding community (and therefore neither was the species ratio) (Fig 2c). However, P22*vir* rose to high abundance and therefore successfully replicated in the community (Supp Fig 4). Additionally, all *S. enterica* isolated from communities exposed to P22*vir* were phage resistant (Table 1).

**Table 1.** Evolution of Resistance and Mucoidy in Phage-Treated Co-Cultures.

| Treatment | # replicate communities | resistant<sup>a</sup> / total isolates | mucoid / total isolates |
| --- | --- | --- | --- |
| ES | 10 | 0/30 | 0/30 |
| ES + P22<sub>vir</sub> | 5 | 15/15 | 0/15 |
| ES + T7 | 5 | 9/9 | 9/9 |

<sup>a</sup> Resistance assayed with cross-streaks on minimal media.

In contrast, *E. coli*-specific T7 attack dramatically altered the final composition of the cross-feeding community. The final frequency of *E. coli* in the community decreased following T7 phage attack, indicating a change in species ratios (p=0.008, Fig 2b). Specifically, *E. coli* density was significantly reduced in the presence of T7 (p = 0.017, Fig 2c). No colonies of *E. coli* could be isolated from the two of the five replicates that were treated with T7 even though low levels of fluorescent protein, and therefore converted species-specific OD, were detected (Fig 2a, Fig 2b). However, all *E. coli* isolates from the remaining three replicate communities were resistant to T7 (Table 1). Surprisingly, growth of *S. enterica* was not constrained by the death of *E. coli*. Instead, *S. enterica* reached between 8- and 35-fold (800-3500%) higher yields in the presence compared to the absence of T7 (Fig2c, Fig 5b). In addition, *S. enterica* started growing before *E. coli*, despite the fact that it depends on *E. coli* acetate excretion for growth (Fig 2a). This led to a rapid increase in biomass in communities with T7 phage (Supp Fig 3). Phage attack on *E. coli* appeared to release *S. enterica* from the constraints typically imposed on community partners when cross-feeding.

### T7-resistant E. coli increase S. enterica densities in the absence of phage

The increase of *S. enterica* in the presence of T7 could result from T7-resistance altering *E. coli* secretion of cross-fed metabolites. To test if T7-resistant *E. coli* secrete more metabolites than sensitive *E. coli*, we assayed the growth of *S. enterica* when paired with *E. coli* isolates in the absence of phage. *S. enterica* reached an average of 1.43-fold (43%) higher density when co-cultured with evolved T7-resistant isolates than with ancestral *E. coli* (Fig 3a, p = 0.004). In these co-cultures, the yield of T7-resistant *E. coli* only reached 67% the yield of the ancestor (Fig 3a, p = 0.004). We also paired P22*vir*-resistant *S. enterica* with ancestral *E. coli* and found that P22*vir* resistance led to a ~2% decrease in *E. coli* yield (Fig. 3b, p < 0.0005), and increased *S. enterica* 5% (1.05-fold) ((Fig 3b, p = 0.012). These results suggest that the divergent impact of T7 and P22*vir* on community composition is in part driven by how resistance to each phage influences host secretion of cross-fed metabolites.

**Figure 3.**
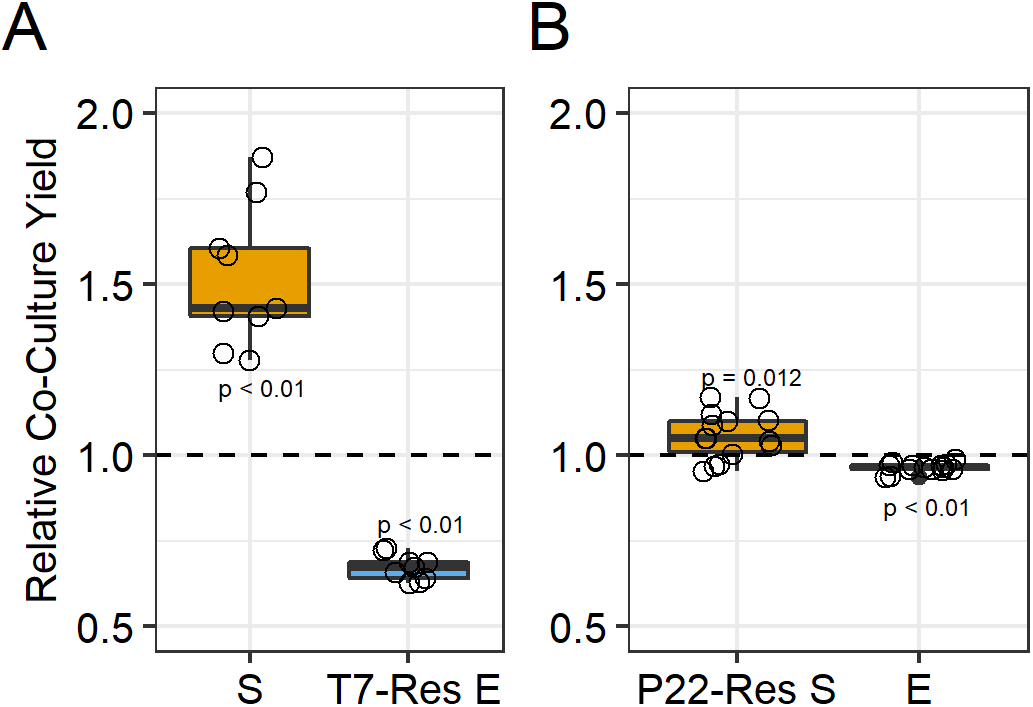
Relative yield of co-cultures with resistant isolates in the absence of phage. The relative yield (CFU) for each strain in co-cultures with one resistant partner as compared to the ancestral co-culture. **A)** Relative yields of co-cultured ancestral *S. enterica* (S) and T7-resistant *E. coli* isolates (T7-Res E) (n = 9). **B)** Relative yields of co-cultured P22*vir*-resistant *S. enterica* isolates (P22-Res S) and ancestral *E. coli* (E) (n = 15). Dashed lines represent standardized yields of ancestral *S. enterica* and *E. coli* co-cultures. Points represent averages of triplicate replicates. Statistical significance determined using Wilcox Sign Test with μ = 1.

### S. enterica grow on cellular debris released by phage lysis

We tested whether consumption of lysed *E. coli* increased the density of *S. enterica* in the presence of T7. When lytic phage burst host cells to release phage progeny, intracellular carbon and nutrients are also released into the environment [21]. To determine if consumption of phage-released cellular debris drove the increase of *S. enterica* during *E. coli*-specific T7 attack, we independently tested *E. coli* and *S. enterica* monoculture growth on cellular debris without phage. We produced cellular debris by sonicating monocultures of *E. coli* and *S. enterica* that had been grown in media with the necessary cross-fed metabolites. Ancestral *E. coli* or *S. enterica* were inoculated at 5×10^5^ cells/ml and grown in lactose minimal medium supplemented with 25% sonication supernatant for 48 hours without phage. We plated for CFUs to determine final yields. While both *E. coli and S. enterica* were able to grow to some extent in lactose minimal media supplemented with cellular debris, we observed different responses. *S. enterica* reached 100-fold higher densities than *E. coli* when both were grown independently on *E. coli* cellular debris (p = 0.050) and 2-fold higher densities than *E. coli* when both were grown independently on *S. enterica* cellular debris (p = 0.046, Fig 4). These results suggest that the consumption of lysed cell components contributes to the increase of *S. enterica* when T7 was added to the cross-feeding community.

**Figure 4.**
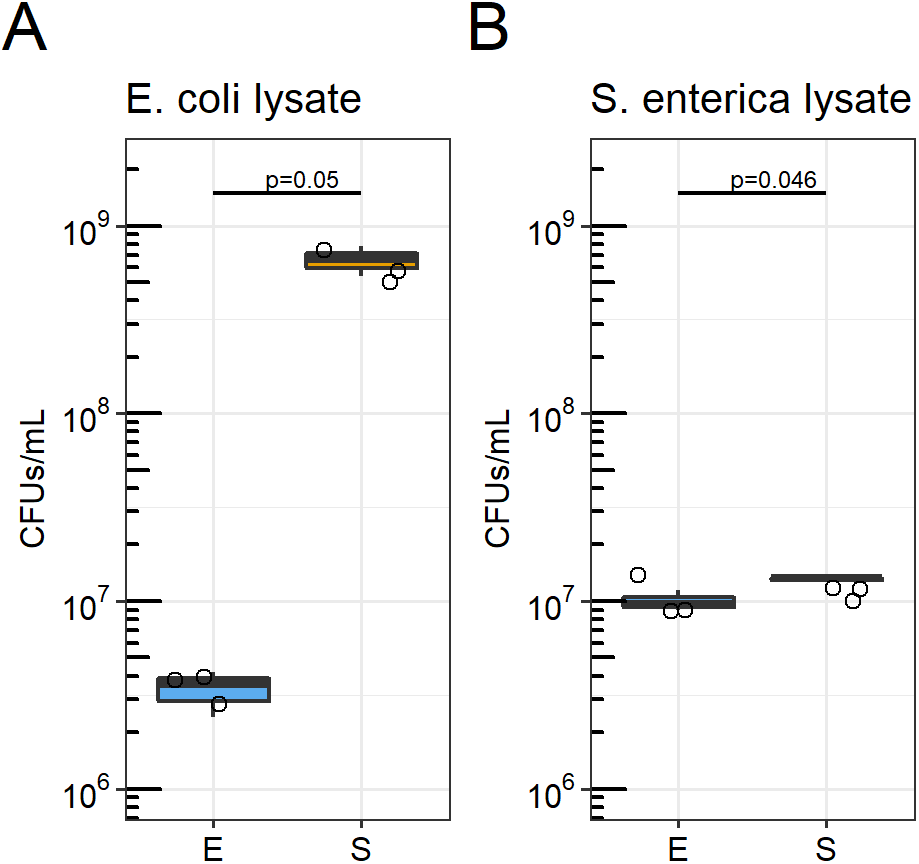
Growth of *S. enterica* and *E. coli* on sonicated cellular debris. *E. coli* (E) and *S. enterica* (S) monocultures were each grown in lactose minimal medium + 25% (v/v) sonicated cellular debris lysate. **A)** Monoculture yields on *E. coli* cellular debris lysate. **B)** Monoculture yield on *S. enterica* cellular debris lysate. Cultures were inoculated with 5 × 10^5^ cells/mL. Statistical significance was tested with a Kruskal Wallis test.

### A modified mathematical model incorporating changes in secretion profiles and cellular debris exchange does not reflect wet-lab experiments

We modified our model to test whether changes in secretion profile and consumption of lysed cells were sufficient to explain the increase in *S. enterica* observed experimentally when T7 was added to the cross-feeding community. Our wet-lab experiments suggested that T7-resistant *E. coli* secrete 50% more acetate than ancestral *E. coli* (Fig 3), and that *S. enterica* grows effectively on cellular debris (Fig 4). We incorporated both these effects into our model. Cellular debris (c_d_) was included as a metabolite that is produced when host cells are killed by phage (Fig 5, Supplemental Methods). As a first approximation, we allowed one lysed host cell to generate enough nutrients to grow one new non-host cell. We found that the increase in acetate production by resistant *E coli* increased *S. enterica* yields 1.9-fold (90%), whereas the incorporation of consumable cellular debris increased *S. enterica* 1.2-fold (20%)compared to the base model with no increase in acetate production or cellular debris exchange. Combining both of these mechanisms increased the final density of *S. enterica* 2.1-fold (210%), well below the ~20-fold (2000%) increase observed experimentally (Fig 5). Therefore, the model suggests that changes in secretion and consumption of lysed cells are insufficient to explain observed patterns, and an additional mechanism likely contributes to *S. enterica* growth when T7 is added to the co-culture.

**Figure 5.**
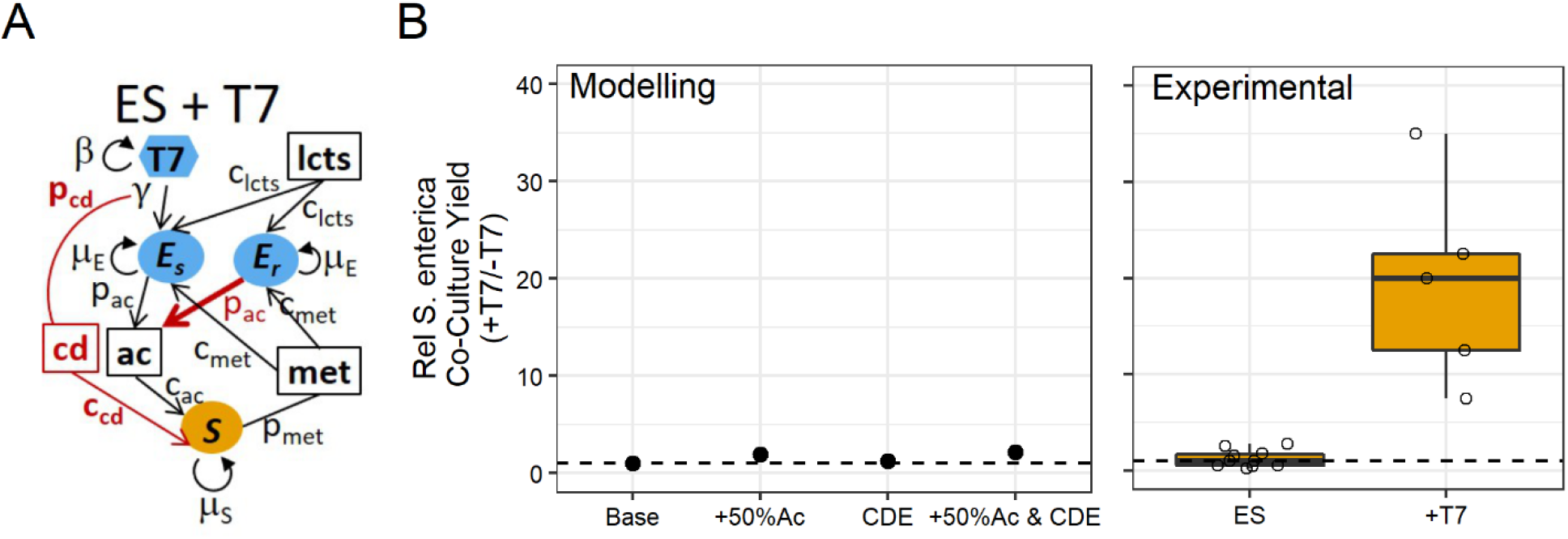
Mathematical model with increased acetate production and cellular-debris exchange increases final densities of *S. enterica* during *E. coli*-specific T7 phage attack. **A)** Schematic of modified model of *E. coli*-specific T7 phage attack. Red text highlights the modifications of cellular-debris (cd) and a 50% increase in acetate production (p_ac_) by T7-resistant *E. coli*. **B)** Relative *S. enterica* co-culture yields of modelling (left panel) and experimental (right panel) results. Results are relative to no phage (-T7) control communities. Base = base model described in figure 1; +50%Ac = T7-resistant *E. coli* produce 50% more acetate compared to T7-sensitive *E. coli;* CDE = cellular debris exchange model where *E. coli* cells lysed by T7 generate cellular debris that can be used by *S. enterica;* +50%Ac & CDE = model combining increase in acetate production when *E. coli* is resistant to T7 phage and consumption of cellular debris by *S. enterica*.

### Evolved partial phage-resistance leads to an increase in S. enterica yields

The phenotype of phage-resistant isolates suggested that partial resistance might contribute to the differential impact of T7 and P22*vir* phage on non-host bacteria. T7 phage-resistant *E. coli* formed mucoid colonies, while P22*vir*-resistant *S. enterica* did not (Fig 6a, Table 1). Mucoidy is frequently associated with partial resistance providing incomplete protection against phage by decreasing efficiency of adsorption [26]. When *E. coli* isolates previously exposed to T7 phage were cross-streaked against T7 phage, no apparent decrease in cell growth was observed, consistent with some level of resistance (Table 1). Additionally, isolates grew in the presence of T7 while ancestral *E. coli* did not (Fig 6b). However, high phage titers were recovered from growth with mucoid *E. coli* isolates; although titers were less than titers recovered from growth on ancestral *E. coli* (Fig 6c). These results suggest that *E. coli* evolved partial resistance may continue to be lysed throughout culturing, increasing the total amount of cell debris available for *S. enterica* to consume in contrast to the full resistance observed in *S. enterica*.

**Figure 6.**
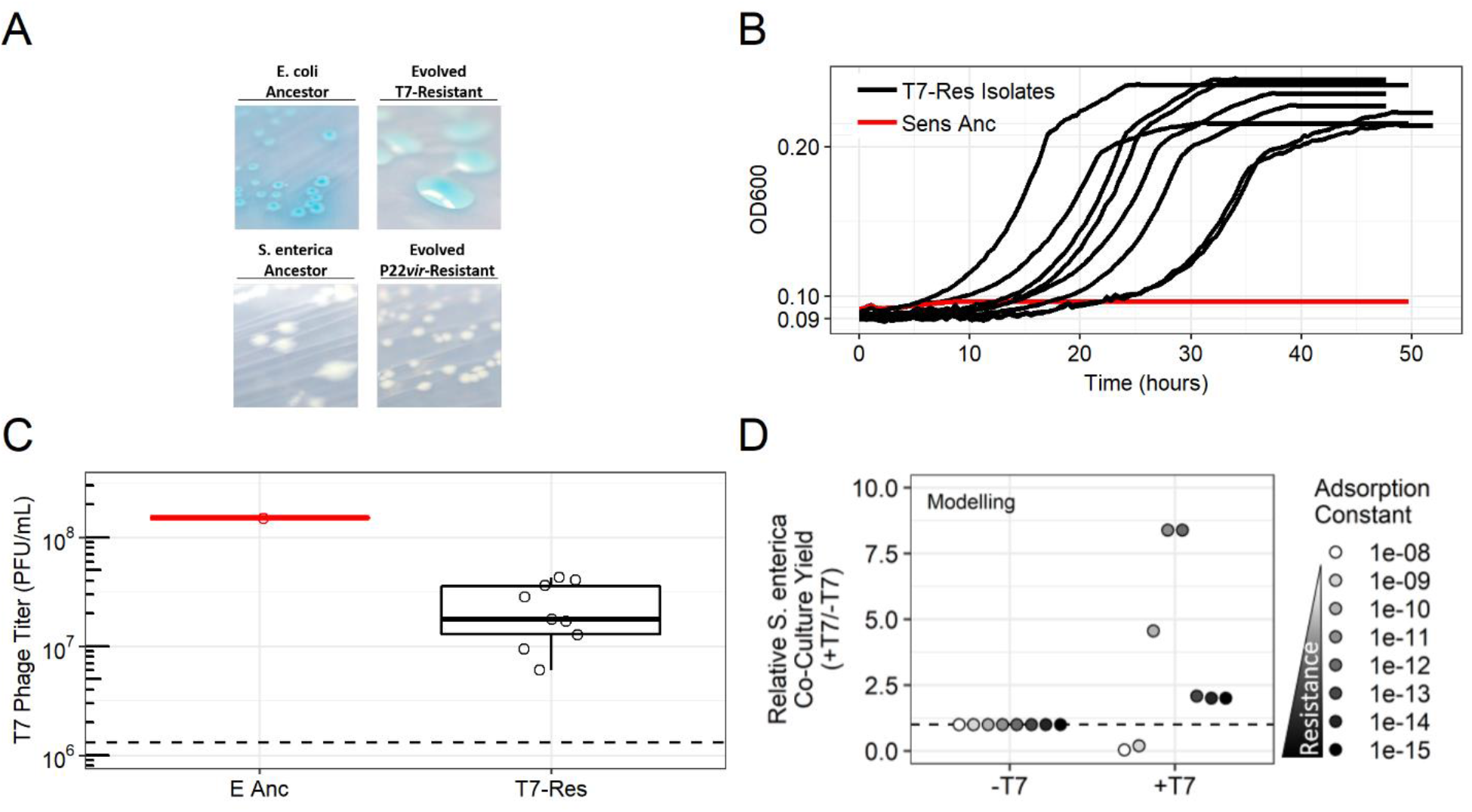
Experimental partial resistance quantification of *E. coli* mucoid T7-resistant and implications on non-host *S. enterica* yield during modeled T7 phage attack. **A)** Morphology comparison of representative *E. coli* T7-resistant isolate or representative P22*vir*-resistant *S. enterica* isolate and ancestral strains on lactose or acetate minimal media, respectively. **B)** Representative growth curves of ancestral *E. coli* (Anc E) or T7-exposed *E. coli* isolates (indicated by isolate number) in the presence of T7 phage. Isolates were grown in lactose minimal medium supplemented with methionine with T7 phage, as indicated (n = 9 isolates). **C)** Titers of T7 phage recovered from infection of *E. coli* isolates. Isolates were inoculated at 0.01 OD and ~10^6^ T7 phage in lactose medium supplemented with methionine in triplicate. After culturing 72 hours in 30°C, phage lysates were made and titered (n = 9 isolates, triplicate). Points are averages of three replicates. E Anc = Ancestral *E. coli*, T7-Res Isolates = mucoid evolved T7-resistant *E. coli* isolates. **D)** Mathematical model with 50% increase in acetate production when *E. coli* is resistant to T7, the ability of *S. enterica* to consume cellular-debris, and partial resistance of *E. coli* against T7 phage. Dashed line = 1.

Genome sequencing also supported partial resistance mechanisms in *E. coli* and a different mechanism in *S. enterica*. We whole-genome sequenced communities treated with either T7 or P22*vir* phage with Illumina sequencing and used *breseq* to identify mutations compared to reference genomes [27]. We focused on mutations unique to each phage treatment and >20% frequency in at least one replicate community. We identified two mutations in the intergenic region of the *clpx* and *lon* genes that reach high frequency in all T7-resistant *E. coli* populations (Table 2). Mutations in the *lon* gene encoding a protease of *E. coli* have been shown to cause mucoid phenotypes in *E. coli* [28]. We also identified a single mutation in four of five *S. enterica* genomes of P22*vir*-treated communities in a Gifsy-1 prophage terminase small subunit that rose to between 80%-90% (Table 2).

**Table 2.** Mutations identified by whole-genome sequencing of phage-treated communities.

| Community Type | Genetic Element | Mutation Location | Mutation | Gene |
| --- | --- | --- | --- | --- |
| ES + P22<sub>vir</sub> | *S. enterica* genome | Coding Region (486/495bp) | Frameshift (+G) | STM2609 - DNA packaging-like protein, small terminase subunit, in *Gifsy-1* prophage |
| ES + T7 | *E. coli* genome | Intergenic (+118/-67) | Δ5 bp :: repeat_region(+) +4 bp | *clpX* - *lon* intergenic region |
|  | *E. coli* genome | Intergenic (+90/-93) | repeat_region (+) +6 bp :: Δ1 bp | *clpX* - *lon* intergenic region |

Finally, we leveraged our model to assess the impact of *E. coli* partial resistance to T7 phage on yield of the non-host, *S. enterica*. We incorporated partial resistance by decreasing the T7 adsorption constant to reduce the frequency of successful phage infection of partially resistant *E. coli* which results in host cells lysing and releasing cellular debris throughout growth. Adding partial resistance, in addition to increasing acetate production of T7-resistant *E. coli* and allowing *S. enterica* to consume *E. coli* cellular debris, led to up to an 8.2-fold (820%) increase in *S. enterica* yield (Fig 6d). Intermediate adsorption values corresponding to partial resistance phenotypes led to the highest yield of *S. enterica* (Fig 6d). These results suggest that multiple mechanisms led T7 attack of *E. coli* to increase the final yield of *S. enterica*.

## Discussion

In summary, our results suggest that lytic phage can dramatically impact communities of cooperating cross-feeding bacteria by changing yields of non-host species. We predicted that attack of a host species with lytic phage would indirectly inhibit cross-feeding species, though resource-explicit models indicated that if resistance evolved, the delay in community growth would be temporary and the community would ultimately reach a similar yield and similar species ratios to communities not attacked by phage (Fig 1). As predicted, attack of *S. enterica* with P22*vir* phage delayed community growth in wet-lab experiments and had little effect on final species ratios (Fig 2, Supp Fig 3). In contrast, *E. coli*-specific T7 attack dramatically altered the final species ratios in favor of *S. enterica*, the non-host, but caused a relatively small delay in community growth (Fig 2, Supp Fig 3). Experimental and mathematical results suggest that a combination of several factors contributed to the increase of *S. enterica* in the presence of T7. These factors include changes in cross-feeding yields, likely from changes in metabolic excretions (Fig 3,5), the ability to grow on cellular debris from phage-lysed cells (Fig 4,5), and a partial resistance mechanism that increased the amount of cellular debris released (Fig 6). These results illustrate that phage can have a wide array of indirect impacts in communities of cooperating bacteria that depend on the metabolic interactions of cooperating cross-feeding partners.

In all cases, we observed that the impact of phage extended beyond the attacked host to all members of the cross-feeding network. P22*vir* inhibited the non-host, while T7 facilitated growth of the non-host. In both cases the effect on growth of non-hosts was indirect and was mediated by changes in the metabolites available in the system. The P22*vir* results demonstrate that cross-feeding metabolic dependencies can make the entire community susceptible to perturbations of a single species. This result mimics previous findings with antibiotic and genetic perturbations in cross-feeding systems [18,19], and highlights the dangers of a cross-feeding lifestyle [29]. In contrast, the indirect effects of T7 were not dependent on cross-feeding. Release of nutrients through cell lysis is likely to generate indirect effects on species abundance independent of microbial interactions. Indeed, viral lysis of bacteria is thought to play a major role in shaping the composition of diverse microbial communities [1,2,4]. Phage are frequently touted as tools for targeted treatment of pathogenic bacteria infections [7,30]. However, our results suggest that even strain-specific phage can have broader impacts on microbial communities, which could lead to diverse phage therapy outcomes.

We have shown two opposing effects of phage attack on a cross-feeding microbial community. One major reason for the divergent effects is likely the identity of the nutrients that each bacterial species needed. In our microbial community, *S. enterica* obtains carbon, in the form of acetate, from *E. coli*, while *E. coli* obtains methionine from *S. enterica*. Bennett et. al. showed that intracellular pools of carbon-compounds are larger than intracellular pools of methionine [31]. Furthermore, it may be easier for cells to scavenge carbon from biomass components than methionine from proteins. Easier access to carbon is supported in our wet-lab experiments, as *S. enterica* reached higher densities on multiple cellular debris types than did *E. coli* (Fig 4). Additionally, the nutritional quality of cellular debris appears to vary, as the difference between *S. enterica* and *E. coli* yields on cellular debris was larger on *E. coli* debris than on *S. enterica* debris (Fig 4). We acknowledge that the metabolites released by phage lysis are likely to differ from those released by sonication of uninfected cells due to phage-mediated host metabolism changes [32]; however, it is unlikely that these differences would qualitatively alter our results. These results suggest that the impact of phage on networks of mutualistic networks may change depending on the identity of cross-fed metabolites and the physiology of lysed cells.

In addition, the magnitude of community responses to T7 phage was influenced by the mechanism of phage resistance that evolved. Both mutations identified in *E. coli* genomes exposed to T7 phage were upstream of the *lon* gene encoding the lon protease, a mutation consistent with a partial resistance phenotype [28]. Down-regulation of the lon protease has been found to cause a mucoid phenotype [28] and negatively regulate the activator of capsular genes, *rscA* [33]. Qimron et. al. showed that knocking out *rscA* also caused mucoidy and partial phage resistance against four phage, including T7 [34]. Our model suggests that partial resistance of *E. coli* significantly increased the indirect effects of *E. coli* – specific T7 phage on *S. enterica*. In the model phage rapidly killed sensitive cells before they rose to 0.2% of the *E. coli* carrying capacity even if we modeled a perfectly efficient conversion of cell debris into biomass (i.e. 1 lysed cell = 1 new cell), the increase of non-host yield was an order of magnitude smaller than observed in experiments with T7. In contrast, partially resistant *E. coli* continued to lyse (though at a lower rate) and generate cell debris. throughout growth. This continual lysis substantially increased yields of the non-host *S. enterica*. Partial resistance should also allow the phage population to be maintained and should therefore lead to lasting changes in species ratios barring further evolution by phage or host bacteria.

Understanding the indirect impacts of phage in microbial communities will be critical as we strive to manage microbial ecosystems. During P22*vir* phage infection, we saw that phage indirectly constrained non-host strains, while T7 phage infection indirectly facilitated the growth of cross-feeders. The qualitative effects and their magnitude were determined by the identity of cross-fed metabolites and consequences of evolved phage-resistance. Identifying how phage influence all bacterial populations in microbial communities is important because of rising enthusiasm for controlling bacteria populations with phage in the food industry and medical field [35,36]. In fact, two independent mouse studies reported that bacteriophage therapy changed abundances of non-host genera in the gut [37,38]. We conclude that understanding nutrient-mediated indirect effects of phage and developing improved methods for inferring cross-feeding mechanism are paramount for predicting the effects of phage attack on relevant and complex microbial communities.

## Material and Methods

### Mathematical Simulations

We used resource-explicit ordinary differential equation models to simulate cooperative growth of *E. coli*, and *S. enterica* (Supplemental Methods). Growth of the bacterial species was governed by Monod kinetics with multiplicative limitation for resources. Production of cross-fed nutrients was growth-dependent. Our base model without phage infection used two equations to directly track *E. coli* (E) and *S. enterica* (S):

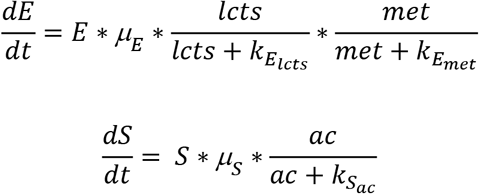

Where E or S is the bacterial population size, μ is a species-specific growth rate (h^−1^), k_x_ is a species- and metabolite-specific Monod constant, and *lcts, met*, and *ac* represent concentrations of lactose, methionine and acetate in g/L.

To the base cooperative cross-feeding model, we used additional equations to incorporate infection by *E. coli*-specific T7 lytic phage or *S. enterica*-specific P22*vir* lytic phage (Fig 1). For example, sensitive *E. coli* (Es) and T7 phage interacted through the following equations:

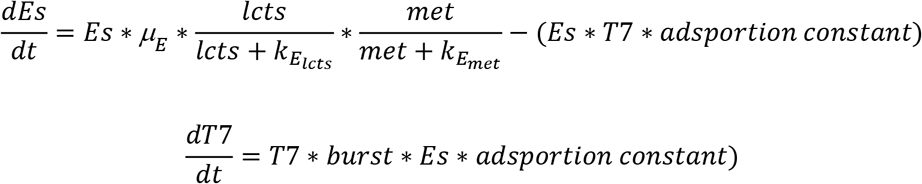

Models contain phage-sensitive (E_s_ or S_s_) host strains, phage-resistant (E_r_ or S_r_) host strains, or non-host (E or S) strains as needed. In a second model, we added an equation for cellular debris (cd). Cellular debris was produced when sensitive host cells were killed by phage. The cellular debris could be consumed by non-host species (Fig 5). We altered acetate production of *E. coli* by changing the production parameters (p_ac_), and encoded partial resistance by changing the adsorption rate parameter (γ). Implicit in our model is the assumption that resistant hosts were present at the beginning of the community growth, and thus resistant genotypes were seeded in at 0.1% of the host population. In practice, resource concentrations were determined algebraically, which improved the stability of the numerical solver. All simulations were run in R with the DeSolve package, using the LSODA solver [39]. Model code is provided in Supplementary Methods and parameters in Supplement Table 1.

### Ancestral Bacterial Co-Culture System and Viral Strains

The ancestral *E. coli* and *S. enterica* strains used in the co-culture system are previously described. Briefly, the ancestral *E. coli* K12 BW25113 *metB::kan* is a methionine auxotroph from the Keio collection with the lac operon reintroduced through conjugation. *S. enterica* LT2 was evolved to secrete methionine [25]. *E. coli* is tagged with a cyan fluorescent protein encoding gene integrated at the lambda attachment site and driven by a constitutive lambda promoter. *S. enterica* is tagged with a yellow fluorescent protein encoding gene under the same promoter and at the same integration site. Co-cultures are performed in lactose hypho minimal medium (5.84 mM lactose, 7.26 mM K_2_HPO_4_, 0.88 mM NaH_2_PO_4_, 1.89 mM [NH_4_]_2_SO_4_, 0.41 mM MgSO_4_)[40]. Monocultures of *E. coli* were supplemented with 80 μM of L-methionine and monocultures of *S. enterica* replaced lactose with acetate. T7 phage is an *E. coli*-specific lytic bacteriophage that cannot attack *S. enterica*. P22*vir* phage is a *S. enterica-specific* lytic bacteriophage that cannot attack *E. coli*. Virus stocks were provided by I. J. Molineaux and were stored at −80°C. Working stocks of phage were grown on ancestral *E. coli* or *S. enterica* cultures grown in minimal medium, as indicated, and stored at 4°C.

### Microbial Community Growth

To assay bacteria community growth, mid-log cultures started from a single bacterial colony were used to inoculate 200μl of medium in a 96-well plate with 10^5^ total cells for each indicated bacterial species per well, and 10^2^ total phage (MOI = 0.01) where indicated. The 96-well plates were incubated in a Tecan Pro200 plate reader for 96-120 hours at 30°C with shaking. OD600, *E. coli*-specific CFP (Ex: 430nm; Em: 490nm), and *S. enterica-specific* YFP (Ex: 500nm; Em: 530nm) fluorescence were read every 20 minutes. After growth in the plate reader, we plated for CFUs of both bacterial species. *E. coli* were selected for on lactose minimal medium plates with excess methionine. *S. enterica* were selected for on citrate minimal medium. X-gal (5-bromo-4-chloro-3-indolyl-β-D-galactopyranoside) in plates further differentiated between *E. coli* and *S. enterica* as *E. coli*-specific β-galactosidase enzymatic activity generated blue colonies, while *S. enterica* colonies remained white. Phage population sizes were measured by plating for plaques on LB with 0.3% LB top agar with a lawn of the ancestral, sensitive host. Phage plates were incubated at 37°C and bacterial plates were incubated at 30°C.

### Testing for Evolution of Phage Resistance: Cross-streaking Assays

Following CFU enumeration, isolated colonies were streaked on minimal media plates for isolation. Resistance to ancestral phage was tested with cross-streaking assays on minimal media plates. Phage stock (~10^8^ PFU) was spread in a line across an LB plate and allowed to dry. Bacterial isolates were then streaked perpendicular to the phage culture. Plates were incubated at 30°C for 24-72 hours. Bacterial isolate streaks with clearing around the phage streak were deemed sensitive and isolates with no clearing were resistant.

### Phenotyping Phage Resistant Isolates

Isolates were cultured alone or in cooperative co-culture as indicated in minimal medium in a 96-well plate. Bacteria were inoculated at 10^5^ cells per well. OD600, and CFP or YFP fluorescence were recorded with the TecanPro200 shaking plate reader for 72 hours. Growth rates were calculated by fitting Baranyi growth curves [41] to fluorescent protein data transformed into OD-equivalents, using experimentally-determined conversion constants, and compared to ancestral strains grown in either monoculture or cooperative co-culture.

### Testing Phage-Mediated Cellular Debris Exchange

*E. coli* was grown in lactose + methionine minimal medium and *S. enterica* was grown in acetate minimal medium. After stationary phase was reached (OD_600_ ~0.5), cells were pelleted and concentrated before being burst with sonication (10, 30s pulses). Sonicated cells were filter sterilized with a 0.22μm filter. Filtered sonication supernatants were checked for sterility by plating. Ancestral bacteria were inoculated at 5×10^5^ cells/mL in lactose minimal medium supplemented with 25% filtered sonication supernatant and incubated at 30°C for 48 hours. Cultures were plated to enumerate CFUs.

### Whole Genome Sequencing of Communities and Analysis

Each community to be sequenced was inoculated from frozen stocks into lactose minimal media and grown at 30°C for 4 days. Total community DNA was isolated using the Zymo Quick-gDNA Miniprep Kit (11-317C). Illumina sequencing libraries were prepared from extracted DNA according to the Nextera XT DNA Library Prep Kit protocol. Prepared libraries were submitted to the University of Minnesota Genomics Center for QC analysis and sequenced on an Illumina Hi-Seq with 125bp paired-end reads. We used the *breseq* tool [27] to align Illumina reads to the following reference genomes: *Escherichia coli str. K-12 substr. MG1655* (Accession No: NC_000913.3), *Salmonella enterica subsp. enterica serovar Typhimurium str. LT2* (Accession No: NC_003197.2, NC_003277.2), Enterobacteria phage T7 (Accession No: NC_001604.1), and Enterobacteria phage P22 (Accession No: NC_002371.2). Mutations lists for resistant populations were filtered such that mutations identified between ancestral strains and reference genomes were removed. We then filtered for mutations that were unique to each phage treatment and focused on mutations that arose in populations to >20% (Table 2).

## Acknowledgements

We would like to thank Ian J. Molineaux (UT-Austin) for the phage isolates; E. Adamowicz, S. Hammarlund, and the Institute of Molecular Virology at the University of Minnesota for constructive conversations; LF was funded by NIH T32 AI83196 and NIH RO1 GM121498-01A1 grants.

## Author Contributions

LF and WRH designed the experiments; LF, JC, and WRH performed mathematical modeling; LF, JA and RHH collected and analyzed wet-lab experimental data; all authors helped write the paper.

