## Supplemental Figures and Methods for "Lytic bacteriophage have diverse indirect effects in a synthetic cross-feeding community" (from another version of this preprint: version 1, 2019-02-25)

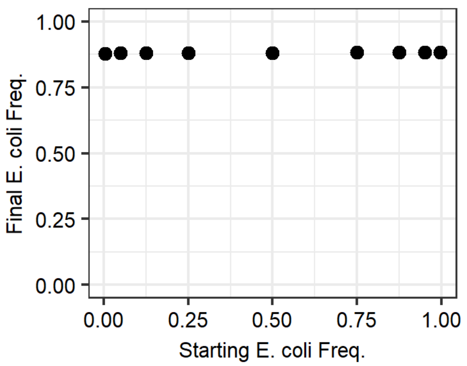


**Supplemental Figure 1. Species frequencies converge in simulated growth regardless of starting frequencies.** We began the simulation with various *E. coli* and *S. enterica* frequencies. Total beginning biomass was 2 x $10^6$ cells regardless of starting ratio. We simulated cooperative growth until all lactose was consumed. Final *E. coli* frequencies of the total bacterial population (E + S) were calculated. Frequencies converged to ~0.88 (88% *E. coli*).

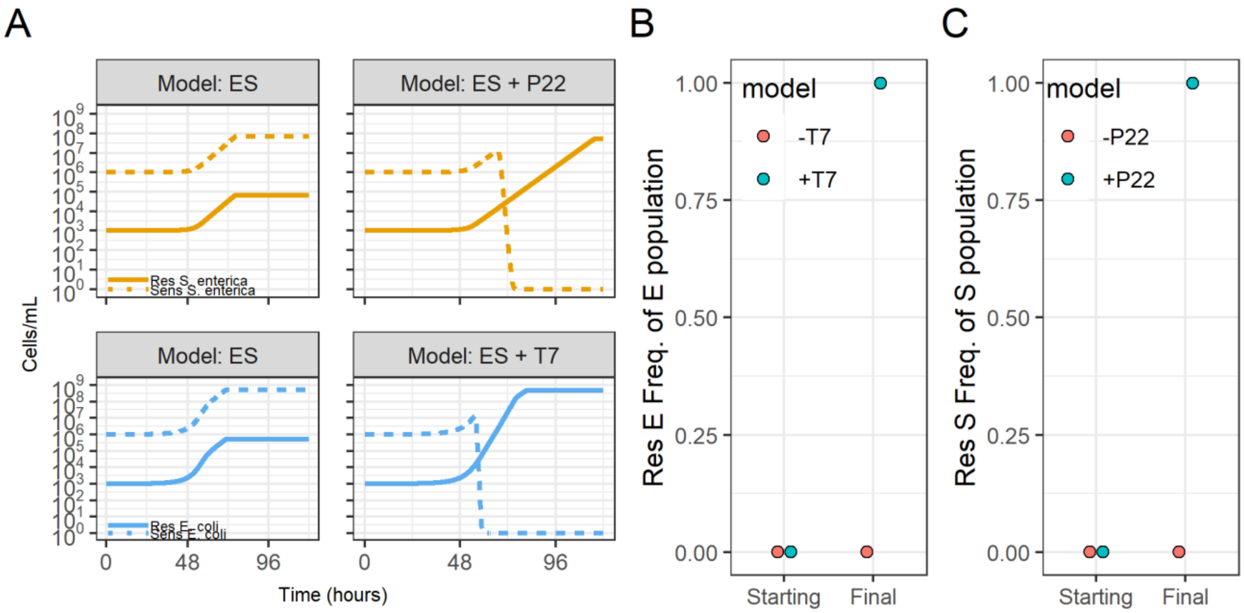


**Supplemental Figure 2. Resistant host cells cannot increase in frequency unless phage are present in the simulated community.** Simulations were run with and without phage. Starting and final frequencies of the host population were calculated for resistant hosts (Er or Sr). While resistant hosts were capable on increasing in absolute density, they did not makes up a significant proportion of the total host populations (sensitive + resistant hosts) unless phage were present. **A)** Growth curves of host simulated growth with and without phage. Blue = *E. coli*, Yellow = *S. enterica*. Solid = resistant genotype, dashed = sensitive genotype. **B)** Frequency of T7-resistant *E. coli* in *E. coli* total population. **C)** Frequency of P22*vir*-resistant in *S. enterica* total population.

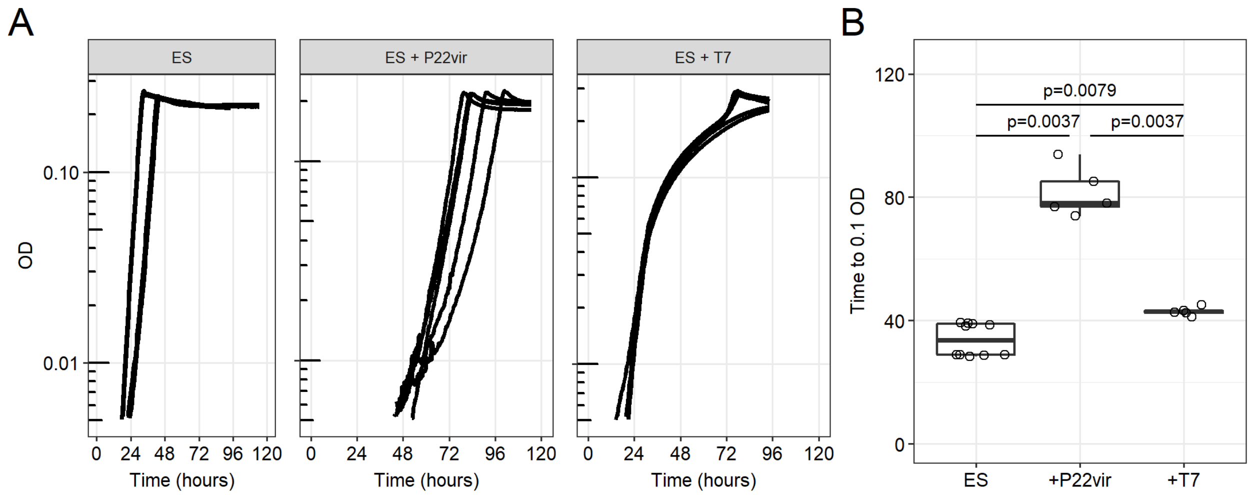


**Supplemental Figure 3. Community OD shows more delayed growth during P22*vir* attack than T7 attack. A)** ODs of cooperating communities treated with the indicated phage were measured every 20min. OD represents a community-level growth because individual species are aggregated. **B)** Summarized time to 0.1 OD for each phage treatment. Statistical significance determined with Mann-Whitney-U test with FDR multiple hypothesis correction.

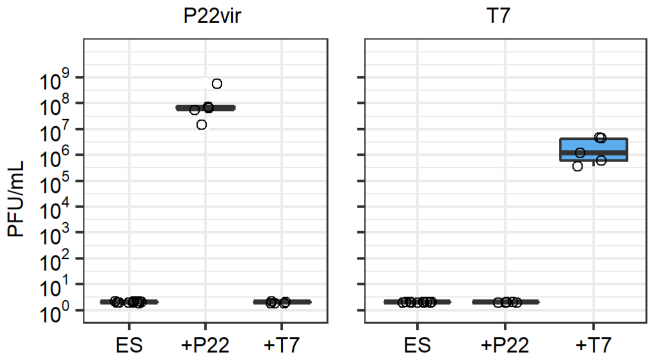


**Supplemental Figure 4. Phage PFU measurements from wet-lab co-culture experiments.** Phage titers of A) P22*vir* and B) T7 were measured for no phage controls (ES, n=10), co-culture with P22*vir* (+P22, n = 5), and co-cultures with T7 (+T7, n = 5).

**Supplemental Table 1. Resource-Explicit Mathematical Model Parameters.**

| Parameter | Description | Value | Units | Reference |
|---|---|---|---|---|
| $\mu_E$ | E, Es, Er growth rate ($hr^{-1}$) | 0.550 (lcts) | $h^{-1}$ | This study |
| $\mu S$ | S, Ss, Sr growth rate ($hr^{-1}$) | 0.156 (ac) | $h^{-1}$ | This study |
| $p_{ac}$ | Es & Er acetate production | $4x\ 10^{-13}$ | g/cell | Harcombe et al 2014 (Adjusted) |
| $p_{met}$ | S methionine production | $1 \times 10^{-12}$ | g/cell | Harcombe et al 2014 (Adjusted) |
| $c_{lcts}$ | Es & Er lactose consumption | $2 \times 10^{-12}$ | g/cell | Harcombe et al 2014 (Adjusted) |
| $c_{ac}$ | S acetate consumption | $3 \times 10^{-12}$ | g/cell | Harcombe et al 2014 (Adjusted) |
| $c_{met}$ | Es & Er methionine consumption | $5 \times 10^{-14}$ | g/cell | Harcombe et al 2014 (Adjusted) |
| $\beta$ | Burst size | 100 | phage progeny/infected cell | De Paepe & Taddei, 2006[a] |
| $\gamma$ | Adsorption rate (successful infection per encounter) | $10^{-8}$ | $min^{-1}$ | De Paepe & Taddei, 2006[a] |
| $k_{lcts}$ | Michaelis-Menton lactose saturation | $7 \times 10^{-7}$ | g/200μl | Harcombe et al 2014 (Adjusted) |
| $k_{ac}$ | Michaelis-Menton acetate saturation | $3 \times 10^{-7}$ | g/200μl | Harcombe et al 2014 (Adjusted) |
| $k_{met}$ | Michaelis-Menton methionine saturation | $3 \times 10^{-7}$ | g/200μl | Harcombe et al 2014 (Adjusted) |

[a] De Paepe & Taddei (2016) used *E. coli* strain MG1655 at 37°C in LB to measure parameters. We decreased our model burst rate ([illegible]) and adsorption rate ($\gamma$) parameters to accommodate slower growth in 30°C growth in lactose minimal media.

**Supplemental Table 2. Measured starting densities and MOIs.**

| Experiment # | Community Type | E (cells/well) | S (cells/well) | Phage (Type) (PFU/well) | MOI |
|---|---|---|---|---|---|
| **1** | ES | $2.25x10^5$ | $1.77x10^5$ | NA | NA |
| **1** | ES + T7 | $2.25x10^5$ | $1.77x10^5$ | $1.6x10^3$ (T7) | 0.007 |
| **2** | ES | $1.1x10^5$ | $1.16x10^4$ | NA | NA |
| **2** | ES + P22*vir* | $1.1x10^5$ | $1.16x10^4$ | $2.1x10^2$ (P22*vir*) | 0.018 |

Supplemental Methods

### An Ordinary Differential Equation Model Exploring Phage Effects on a Cross-Feeding Microbial Co-culture Community in R

**Lisa Fazzino[1,4], Jeremy Anisman[2,4], Jeremy M. Chacón[3,4], Richard H. Heineman[5], William R. Harcombe*[1,3,4]**

[1]*Department of Microbiology & Immunology, University of Minnesota, Minneapolis, MN;*

[2]*College of Continuing & Professional Studies, University of Minnesota, Minneapolis, MN;*

[3]*Ecology, Evolution, and Behavior, University of Minnesota, Minneapolis;*

[4]*Biotechnology Institute, University of Minnesota, Minneapolis;*

[5]*Biology Department, Kutztown University, Kutztown, PA*

#### 1. Resource-Explicit ODE Model of Phage Infection of a Bipartite Cross-Feeding Microbial Community

We used the following systems of ordinary differential equations to model changes in biotic populations of an engineered mutualistic co-culture community comprised of *Escherichia coli* (E) and *Salmonella enterica* (S).[1] We added equations to model the effects of E-specific phage (T7) and S-specific phage (P22*vir* or P22). Bacterial populations were comprised of phage-sensitive (s) and phage-resistant (r) individuals, as indicated by a lowercase letter. Because nutrient concentrations are conserved in the system, they are modeled using algebraic equations because we have found that this improves the performance of the numerical solver.[2] The model used the units of cells/mL and g/µL for nutrient resources. Definitions of model parameters are in Table S1. Parameters include interactions between biotic populations, phage population parameters, nutrient uptake, and conversion parameters. Although we used parameter values and variable names that mimic the *E. coli / S. enterica* wet-lab community, these models are generalizable to any species and phage system.

##### 1.1 Base Monod Growth Model Without Phage Infection

We started with the following set of equations as a base model of *E. coli* and *S. enterica* cross-feeding interactions in lactose minimal medium:

Biotic Equations:

$$\frac{dEs}{dt} = (Es)(\mu_{Es})\left(\frac{lcts}{(lcts + k_{Elcts}}\right)\left(\frac{met}{(met + k_{Emet}}\right)$$

$$\frac{dS}{dt} = (S)(\mu_S)\left(\frac{ac}{(ac + k_{Sac}}\right)$$

Abiotic Equations:

$$lcts_{t=t} = lcts_{t=0} - (Es_{t=t} - Es_{t=0})(c_{Elcts})$$

$$met_{t=t} = met_{t=0} - (Es_{t=t} - Es_{t=0})(c_{Emet}) + (S_{t=t} - S_{t=0}) * p_{Smet}$$

$$ac_{t=t} = ac_{t=0} + (Es_{t=t} - Es_{t=0})(p_{Eac}) - (S_{t=t} - S_{t=0}) * c_{Sac}$$

where $\mu_x$ represent isolate-specific growth rates. In this model formulation, *E. coli* growth is limited by lactose (lcts) and methionine (met) concentrations in a multiplicative manner, and saturates with Monod half-saturation parameter ($k_x$). Accumulation of produced metabolites is growth-dependent.

1.2 Modelling Phage Infection of Cross-Feeding Bacterial Community

We added phage infection of either host species to the model equations by including a phage equation, a resistant host equation with a different maximum growth rate parameter (μEr), and a term in the sensitive host to represent phage-mediated death. Phage-mediated death is modeled using a classic, linear, host-parasite interaction. However, since T7 phage reproduction significantly decreases on stationary phase cells[3], in our model, T7 phage only produced new progeny from actively growing hosts. Abiotic equations were also changed to accurately represent metabolite consumption and production by additional bacterial genotypes.

The following example set of equations model T7 phage infection of *E. coli:*

Biotic Equations:

$$\frac{dEs}{dt} = (Es)(\mu Es)\left(\frac{lcts}{lcts + k_{Elcts}}\right)\left(\frac{met}{(met + k_{Emet}}\right) - Es\,(T7)(\gamma)$$

$$\frac{dT7}{dt} = (T7)(\beta)(Es)(\mu Es)\left(\frac{lcts}{lcts + k_{Elcts}}\right)\left(\frac{met}{(met + k_{Emet}}\right)(\gamma)$$

$$\frac{dEr}{dt} = (Er)(\mu Er)\left(\frac{lcts}{lcts + k_{Elcts}}\right)\left(\frac{met}{(met + k_{Emet}}\right)$$

$$\frac{dS}{dt} = (S)(\mu S)\left(\frac{ac}{(ac + k_{Sac}}\right)$$

Abiotic Equations:

$$lcts_{t=t} = lcts_{t=0} - (Es_{t=t} + Er_{t=t} - Es_{t=0} - Er_{t=0})(c_{Elcts})$$

$$met_{t=t} = met_{t=0} - (E_{t=t} + Er_{t=t} + (Es_{t=t} * T7_{t=t} * \gamma) - Es_{t=0} - Er_{t=0})(c_{Emet}) + (S_{t=t} - S_{t=0}) * p_{Smet}$$

$$ac_{t=t} = ac_{t=0} + (Es_{t=t} + Er_{t=t} + (Es_{t=t} * T7_{t=t} * \gamma) - Es_{t=0} - Er_{t=0})(p_{Eac}) - (S_{t=t} - S_{t=0}) * c_{Sac}$$

where $Es\ (T7)(\gamma)$ represents the number of sensitive *E. coli* host cells killed by T7 phage infection with the adsorption constant ($\gamma$), β is the phage burst size, and Er represents the phage-resistance genotype that is seeded in at 0.1% of the sensitive population level.

To model P22*vir* infection of *S. enterica*, similar equations as above were used with Ss, Sr, and P22 equations to replace Es, Er, and T7.

#### 2. Resource-Explicit ODE Model Including Phage-Lysis Mediated Exchange of Cellular Debris

To our base model, we added the ability of non-host cells to consume cellular debris released during phage lysis of host cells. To accomplish this, we added a metabolite equation to represent cellular debris (cd). Production of cellular debris occurred when phage hosts died by phage lysis, with production rate parameter p_e_cd. Non-hosts (S) could grow using cellular debris with Monod kinetics. The consumption parameter c_s_cd describes the amount of cellular debris required to produce an *S. enterica* cell from cellular debris. *S. enterica* cells growing on acetate and cellular debris could grow faster than *S. enterica* cells growing on either resource alone because acetate and cellular debris could be used as substitutes for each other. Furthermore, growth on both substrates was additive.

##### 2.1 Attacking *E. coli* with T7 phage allowing *S. enterica* to consume lysed *E. coli* cellular debris.

To add the consumption of cellular debris (cd), we modified the previous set of equations describing phage infection of the host to include a cellular debris metabolite equation, and added a growth term to the non-host species to describe growth of cellular debris. Other metabolite equations were modified to include consumption of metabolites by cells that had been lysed during that time step.

$$cd = Es\ (T7)(\gamma)(p_{e_{cd}}) - (Scd * c_{s_{cd}})$$

$$\frac{dS}{dt} = (S)(\mu S)\left(\frac{ac}{(ac + k_{Sac}}\right) + (S)(\mu S)\left(\frac{cd}{(cd + k_{Scd}}\right)$$

#### 3. Model Parameters

Table of model parameters with references can be found in Table S1.

#### 4. Simulation Framework

ODE models were numerically integrated using the lsoda solver from the deSolve package in R. Time was in units of hours. We set hmax = 0.001, rtol = $1\times10^{-10}$ (default), and atol = $1\times10^{-13}$, which improved the solutions. Code to run a sample model with a subset of parameters is included in the supplementary R file.

```
# Resource-explicit mathematical models for Fazzino et al. 2019. BioRxiv
# R Script File
# Set working directory with setwd()

library(ggplot2)
library(dplyr)
library(tidyr)
library(deSolve)

# T7-infection of cross-feeding E. coli
coop_e <- function(t, n, parameters){
  with(as.list(c(t, n, parameters)),{

    lcts = lcts_0 - (Es + Er + Edead - Es_0 - Er_0) * c_e_lcts
    met = met_0 - (Es + Er + Edead - Es_0 - Er_0) * c_e_met + (S - S_0) * p_s_met
    ac = (ac_0 + ((Es + Er + Edead - Es_0 - Er_0) * p_e_ac) - ((S - S_0) * c_s_ac))

    gEs = Es * mu_es * (lcts / (lcts + k_e_lcts)) * (met / (met + k_e_met))
    dEs = gEs - (Es*T7*1e-8) #1e-8 = adsorption rate
    dEdead = Es*T7*1e-8 #1e-8 = adsorption rate
    dEr = Er * (mu_er * (lcts / (lcts + k_e_lcts)) * (met / (met + k_e_met)))

    dS  = S * mu_s * (ac / (ac + k_s_ac))

    dT7 = T7 * burst * gEs * 1e-8 #1e-8 = adsorption rate

    return(list(c(dEs, dEr, dS, dT7, dEdead)))
  })
}

parameters = c(mu_es = 0.55,        # /hour grown on lactose
        mu_er = 0.55,        # /hour grown on lactose
        k_e_lcts = 7e-7,     # g/200microliters
        k_e_met = 3e-7,      # g/200microliters
        c_e_lcts = 2e-12,    # grams lcts used per E cell made
        c_e_met = 5e-14,     # grams met used per E cell made
        p_e_ac = 4e-13,      # grams ac produced per E cell made

        mu_s = 0.17,         # /hour grown on ac
        k_s_ac = 3e-7,       # gram/200microliters
        c_s_ac = 3e-12,      # grams ac used per S cell made
        p_s_met = 1e-12,     # grams met produced per S cell made
        burst = 100,         # T7 phage produced per lysed infected Es cell

        lcts_0 = 1e-3,       # grams/200microliters
        met_0 = 1e-10,       # grams/200microliters
```

```
            ac_0 = 0,
            glc_0 = 0,

            Edead_0 = 0,        # cells/200microliters
            Es_0 = 1e6,         # cells/200microliters
            Er_0 = 1e3,         # cells/200microliters
            S_0 = 1e6)          # cells/200microliters

start_density = c(Es = 1e6, Er = 1e3, S = 1e6, T7 = 1e2, Edead = 0)
time_to_sim = seq(from = 0 , to = 125, by = 0.1)
out = ode(y = start_density,
         times = time_to_sim,
         func = coop_e,
         parms = parameters,
         rtol = 1e-14,
         hmax = 0.01)
plot(out)
```
